# Modeling Sensorimotor Processing with Physics-Informed Neural Networks

**DOI:** 10.1101/2024.09.14.613030

**Authors:** Adriana Perez Rotondo, Alessandro Marin Vargas, Michael Dimitriou, Alexander Mathis

## Abstract

Proprioception is essential for planning and executing precise movements. Muscle spindles, the key mechanoreceptors for proprioception, are the principle sensory neurons enabling this process. Emerging evidence suggests spindles act as adaptable processors, modulated by gamma motor neurons to meet task demands. Yet, the specifics of this modulation remain unknown. Here, we present a novel, physics-informed neural network model that integrates biomechanics and neural dynamics to capture spindle function with high fidelity and efficiency, while maintaining computational tractability. Through validation across multiple experimental datasets and species, our model not only outperforms existing approaches but also reveals key drivers of variability in spindle responses, offering new insights into proprioceptive mechanisms.

## Introduction

Deep learning has driven the development of powerful models of brain functions that excel at explaining neural activity (1–10). While deep learning provides highly effective function approximators (11, 12), these models suffer from a major drawback: they lack interpretability. One common strategy to address this issue is the application of interpretable AI techniques (13–16). Another promising approach is to incorporate physics and biological principles directly into the model architecture and training objective, offering both interpretability and alignment with the system’s dynamics (17–20). In this work, we develop such a framework, integrating deep learning with mechanistic principles to develop a framework for modeling the sensorimotor system.

Proprioception plays a pivotal role in sensorimotor processing. Muscle spindles are the principal receptors for proprioception and are considered to be the most complex sensory organs outside of the special senses (21). Despite over six decades of dedicated research on muscle spindles, obtaining comprehensive data simultaneously encompassing fusimotor and afferent components during both active and passive naturalistic movements across diverse tasks and animal species remains a significant challenge (22–24). Modeling muscle spindles is essential not only for understanding the mechanisms by which muscle states are converted into neural afferent firing rates but also for elucidating how these afferents contribute to sensory-motor integration, motor learning, and postural control (21, 24, 25). Various models of muscle spindles have been developed to fill the knowledge gap in experimental recordings. These models range from abstract (statistical) models (26–30) to biophysicaly detailed models (31–37). The former lack an understanding of the underlying physical variables, making generalization across datasets challenging, while the latter, due to their computational complexity, pose integration difficulties into broader models.

In this study, we leverage physics-informed neural networks (PINNs) (17, 19, 38) to present a novel model of muscle spindles that bridges the gap between physical relevance and ease of training with integration into broader models. The unique advantage of PINNs lies in their ability to incorporate domain knowledge, thereby enhancing model interpretability and accuracy. We successfully fit single-trial afferent responses from three different mammals in both passive and active conditions. Our model outperforms both statistical and classical mechanistic models in capturing the wide variety of data, seamlessly combining the flexibility of data-driven approaches with the interpretability of mechanistic models. We uncover variations in afferent dynamics, highlighting inherent differences in muscle spindle tuning across animals and between active and passive movements. Notably, our model allows us to map these variations to specific physical parameters, providing valuable insights into the complex relationship between fusimotor drive, sensory encoding and physiological factors. This study represents a significant step towards a more comprehensive understanding of proprioceptive mechanisms and their intricate interplay with diverse physiological variables.

## Results

### Spindle Modeling Background

To understand the challenges with modeling muscle spindles, it is critical to review the key components. Muscle spindles are sensory organs embedded within most vertebrate skeletal muscles. As the muscle contracts and stretches, so does the muscle spindle (24, 25, 40). Unlike the surrounding extrafusal muscle fibers responsible for movement generation, muscle spindles contain specialized intrafusal muscle fibers that change length with the extrafusal muscle fibers (41, 42). These intrafusal fibers are innervated by primary and secondary sensory endings, which transduce the mechanical state (tension) of the intrafusal fibers into electrical signals (receptor potentials (21)). Receptor potentials can then trigger action potentials in the sensory neuron that travel along afferent pathways to the central nervous system, propagating information on the state of the muscle.

Thus, the relationship between changes in joint angle and sensory afferent firing rates involves three key processes (Figure 1A). First, a joint movement translates into a change in the length of the muscle fascicles where the muscle spindle is located (43–47). Second, *mechanical filtering* within the muscle transforms this change in fascicle length into a change in tension within the intrafusal muscle fibers (31, 44, 45). Importantly, fusimotor inputs can independently modulate this tension (43, 46, 48, 49) In fact, *γ* motor neurons innervate intrafusal fibers and their activation influences tension independently of changes in muscle length (25). Finally, *mechanotransduction* converts this tension change into a receptor potential within the sensory endings of the spindle organs, ultimately leading to a change in their firing rate (50–53).

**Figure 1.**
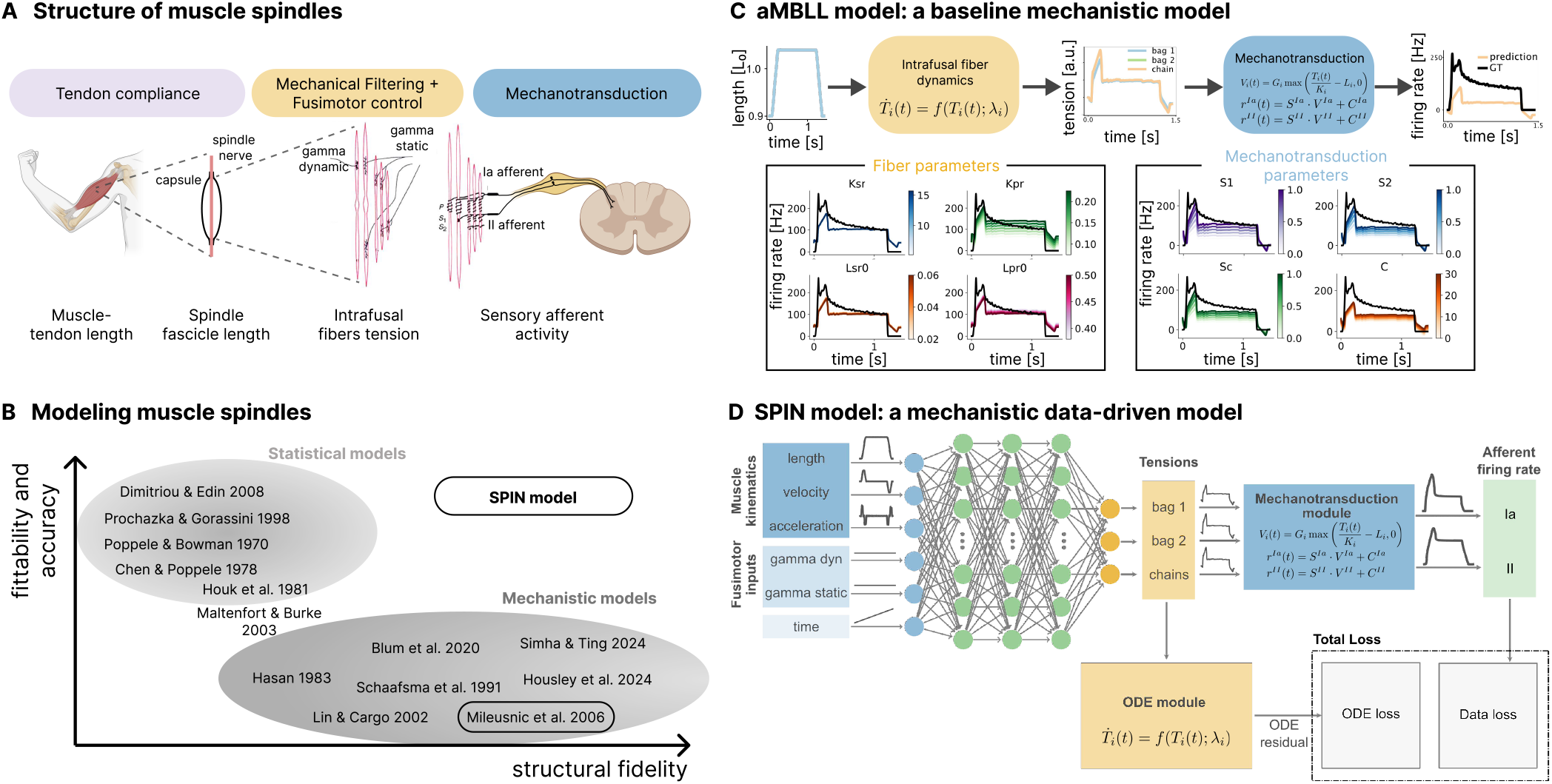
Modeling muscle spindles with PINNs. **A** The three key processes that transform joint movement into sensory afferent activity. Figure adapted from (39) and BioRender. **B** Various models of muscle spindles have been proposed, falling into two main categories: statistical and mechanistic. Statistical models lack structural fidelity but are computationally efficient, while mechanistic models are based on the structural and functional understanding of spindles but are hard to train and computationally inefficient. The proposed SPIN model combines these approaches, providing a mechanistic model that is easy to train on experimental data. **C** The mechanistic components of SPIN are based on an adapted Mileusnic-Brown-Lan-Loeb (aMBLL) model (34). The top of the panel shows a schematic representation of the model, which transforms muscle stretch (and fusimotor activity) into intrafusal fiber tensions using an anatomically motivated differential equation. These tensions are then combined to produce the Ia and II afferent firing rates. This model, with its 48 parameters tuned for cat *soleus* muscle spindle properties, struggles to fit recordings from the cat *triceps surae* even when the model parameters are adjusted (bottom panel). **D** The SPIN model estimates the intrafusal fiber tensions with a multi-layer perceptron that takes muscle kinematics (normalized muscle length, velocity, and acceleration), fusimotor inputs (*γ* static and dynamic activity), and time as inputs. The intrafusal fiber tensions are transformed into Ia and II firing rates based on the same equations as the aMBLL model. We train the model using a loss function that minimizes the error between the experimental and predicted firing rates (Data loss) and constrains the predicted tensions to satisfy the aMBLL defined differential equation (ODE loss).

To better understand their underlying mechanisms, various models of muscle spindles have been proposed. These can be divided into two categories: statistical and mechanistic (Figure 1B). Statistical approaches to modeling spindles often use linear transfer functions (26, 27) or simple nonlinear functions (29, 30) to describe how muscle kinematics (length and velocity) affect afferent firing rate. For instance, some models have proposed linear or pseudo-linear (linear with half-wave rectifiers) representations with either muscle kinematics inputs (26, 27, 54) or muscle force and yank inputs (55). These models can represent the ensemble activity of a population of afferents for a specific range of movements (29, 56). However, they often struggle to generalize to the response of individual afferents for a wide range of motions and behaviors. Crucially, they fail to account for the underlying anatomical, structural, and physiological mechanisms of spindle function.

In contrast, mechanistic models provide a more complete, mathematical representation of spindle afferent responses based on the understanding of the structure and the function of spindles. Variations exist in the level of anatomical detail included in each model (see a comprehensive review by (56)). Mechanistic models typically incorporate detailed models of intrafusal fibers (31–34). This often involves differential equations describing the tension of each intrafusal fiber type based on length changes and fusimotor inputs.

Although mechanistic models excel in linking muscle spindle response directly to its anatomical and functional components in an interpretable manner, they face two key limitations. Firstly, solving the differential equations makes them computationally expensive for simulations. Secondly, increased model detail necessitates tuning numerous parameters (over 30) to match afferent responses. These parameters are often based on limited anatomical data or sparse recordings of single muscles in cat experiments, restricting their generalizability. Consequently, existing mechanistic models have rarely been used to predict single afferent responses beyond a single animal species or recording technique (31, 33, 34).

### Spindle physics informed neural networks

We propose a novel approach to muscle spindle modeling using physics-informed neural networks. In essence, our Spindle Physics Informed Neural network (SPIN) model allows one to learn a model of muscle spindles based on recordings (data loss) and known dynamics (ODE loss) (Figure 1C,D). In the following we will explain these components.

The ODE-part is based on the classic mechanistic models of muscle spindles (31–34). In particular, Mileusnic et al. (34) construct a physiologically realistic model of the spindle with differential equations that capture the dynamics of anatomical components found in muscle spindles. We refer to it by the acronym of the four authors: MBLL for MileusnicBrown-Lan-Loeb.

The MBLL model defines a single differential equation with different parameters to capture the three types of intrafusal fiber dynamics: bag 1, bag 2 and chain. The solution of the equation gives a tension for each fiber type under a given fascicle stretch (length, velocity, and acceleration) and fusimotor input (constant activation of static or dynamic *γ* motor neuron). Following anatomical observations, bag 1 fiber is the only fiber that receives inputs from dynamic fusimotor efferent endings and bag 2 and chain fibers receive static fusimotor inputs. The tensions of the three fibers are then combined to obtain the primary Ia and secondary II afferent activity. The Ia afferent activity has contributions from all three fiber types whereas the II afferent activity is only determined by the bag 2 and chain fiber tensions.

We adapt the MBLL model and use it as SPIN’s mechanistic model, referring to it as the adapted Mileusnic-BrownLan-Loeb (aMBLL) model (Figure 1C). This model forms the foundation for the SPIN model described later. Key modifications to the original MBLL model include removing the mass term in the intrafusal fiber dynamic equation, reducing it from a second-order to a first-order differential equation, and simplifying the number of fiber parameters. Additionally, we revised the mapping of fiber tensions to afferent firing rates by expressing the firing rate as a linear combination of the afferent ending potentials (see Methods).

The SPIN model, similar to the aMBLL model, focuses on representing the tension of the intrafusal fibers (Figure 1D). A conventional feed-forward neural network with three hidden layers (each containing 128 units) takes six inputs (fascicle length *l*, velocity *v*, acceleration *a*, dynamic *γ* activation *γ*_*d*_, static *γ* activation *γ*_*s*_, and time *t*) and outputs the tensions of the three fiber types. Thus, the predicted fiber tensions are defined by the neural network transformation

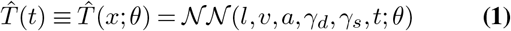

where 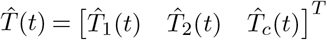 is the vector with tensions of all three fiber types, *θ* denote the weights and biases of the neural network, *x* is the vector with all the inputs to the model—the fascicle length *l*, velocity *v* and acceleration *a*, the fusimotor inputs *γ*_*d*_ and *γ*_*s*_ and time *t* (see Methods).

These tensions are then transformed into Ia and II afferent firing rates using pre-defined equations referred to as the mechanotransduction or output module. The afferent firing rates are defined as:

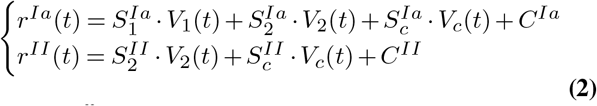

where 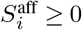 are nonnegative scalars representing the contribution of fiber type *i* to the activity of the afferent aff ∈ {*Ia, II*}. The constant *C*^aff^ ≥ 0 represents the baseline firing rate of the afferent aff. *V*_*i*_ represent the receptor potential of fiber type *i* defined as follows:

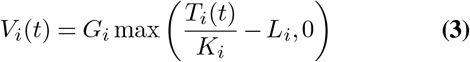

where *K*_*i*_ denotes the sensory ending constant, which quantifies how a change in fiber tension relates to a change in ending stretch. The threshold length, *L*_*i*_, is the sensory ending stretch beyond which ion channels open, causing depolarization in the sensory ending. The scaling constant *G*_*i*_ relates the stretch to the afferent potential.

The overall output model comprises 16 tunable parameters, denoted as *λ*_*m*_. These parameters (or combinations of them) can be related to specific anatomical and physiological mechanisms. Specifically, they account for the contributions of different fiber types to various afferent responses, differences in sensitivity to sensory ending stretches, variability in baseline firing rates, and partial occlusion (34, 57). We also illustrate how variations of some of these parameters influence afferent firing (Figure 1C).

We define an overall muscle spindle model that incorporates muscle kinematics, fusimotor inputs, and Ia and II afferent activity outputs. The parameters *θ* and *λ*_*m*_ determine the model output. The model is trained through supervised learning, where the weights of the neural network *θ* and the parameters of the mechanotransduction module *λ*_*m*_ are optimized through backpropagation of a loss function derived from measurements (labeled training data).

The essence of PINNs is that they utilize information about the system’s dynamics as well as data to train the model (Figure 1D). This information is integrated into the model by adding terms to the loss function that constrain some of the system’s outputs to satisfy specific constraints. We denote these additional losses as the ordinary differential equation (ODE) loss ℒ_*ODE*_ and the initial condition loss ℒ_*IC*_. The total optimization loss is therefore expressed as:

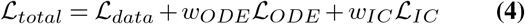

Here *w*_*ODE*_ and *w*_*IC*_ are hyperparameters that weight the contribution of each loss function to the total loss function. For the data loss, we use the classic mean square error (MSE) loss on the firing rate. Typically, we train the model independently for each afferent (Ia and II), thus:

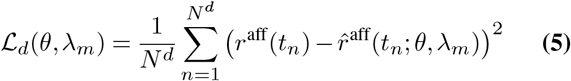

where *r*^aff^(*t*_*n*_) is the measured afferent firing rate at time *t*_*n*_ and 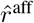 is the model prediction and *N* ^*d*^ is the number of measurements of the afferent firing rate (trials times timepoints per trial).

The initial condition loss penalizes the model when its predicted tensions 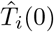 differ from a target initial tension *T*_*i*_(0)

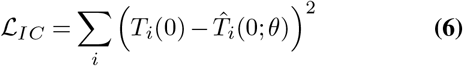

where the sum is over the three tension types: bag 1, bag 2, and chain. The crucial biomechanical knowledge is captured by the ODE loss:

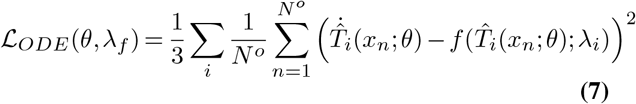

where 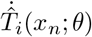 denotes the time derivative of the tension prediction 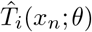. This loss function constrains the tensions predicted by the model to satisfy the dynamics defined by the function 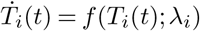. *f* based on the differential equations of the aMBLL model Eq. (17). Although *f* has the same form for each fiber type *i*, it has different parameters *λ*_*i*_. We refer to the ensemble of parameters *λ*_*i*_ for each fiber type as the *fiber parameters*: *λ*_*f*_ = *λ*_*bag*1_ ∪*λ*_*bag*2_ ∪*λ*_*chain*_.

Overall, the SPIN model is fully defined by the fiber parameters (*λ*_*f*_), mechanotransduction parameters (*λ*_*m*_), and the neural network weights (*θ*). The parameters *λ*_*f*_ and *λ*_*m*_ represent physiological properties of the muscle spindle (which are measurable), while the neural network weights (*θ*) are only responsible for generating intrafusal fiber tensions that satisfy the known dynamic equations. Note that the mechanotransduction parameters contain the physiology knowledge, but this part is not encoded as a differential equation.

SPIN can be regarded as a mechanistic data-driven model. If the values of the model parameters are known from prior experimental studies, SPIN can be trained without the need for new experimental data—using only the ODE loss (ℒ_*ODE*_)—to create an efficient surrogate for the mechanistic aMBLL model. On the other hand, if we have experimental recordings from afferents but *incomplete* knowledge of the model parameters, SPIN can be trained by optimizing both the neural network weights (*θ*) and a subset of the model parameters (*λ*_*f*_ and *λ*_*m*_) to minimize the total loss (ℒ_*total*_). This results in a mechanistic model of the muscle spindle that aligns with both the general anatomical and physiological properties of muscle spindles, while also fitting the specific experimental data from the recorded spindle.

### Model identification

Can SPIN accurately learn a mechanistic model from firing rate measurements?

To address this question, we trained SPIN using synthetic data generated from the aMBLL model with known ground truth parameters, 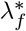 and 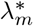, by solving its differential equations directly. Specifically, we generated muscle stretch profiles, used them as inputs to the aMBLL model, and obtained both predicted firing rates and fiber tensions. These synthetic data were then used to train SPIN in the same manner as we would with experimental data. This allowed us to evaluate not only the model’s fit to the firing rate but also its ability to capture the underlying mechanistic features of the system (see Methods).

We used classic ramp-hold-release muscle stretches and triangular stretches at various speeds, generating a total of 100 stretch trials with trial duration ranging from 1.5 to 3 seconds (Figure 2A). To simulate potential noise in experimental recordings, we added multiplicative Gaussian noise to the firing rate data generated by the aMBLL model (see Methods). The dataset was split into five folds using an 70/10/20 train/val/test split, with the SPIN model trained on 70% of the data and tested on unseen trials. We compared the performance of SPIN against several baseline models, including a multi-layer perceptron (MLP) with the same architecture as SPIN’s neural network, and a pseudo-linear model (a linear model with a half-wave rectifier activation function) that used muscle kinematics inputs (length, velocity, and acceleration) (see Methods). Note that the MLP also differs in terms of the constraining mechanotransduction module from SPIN.

**Figure 2.**
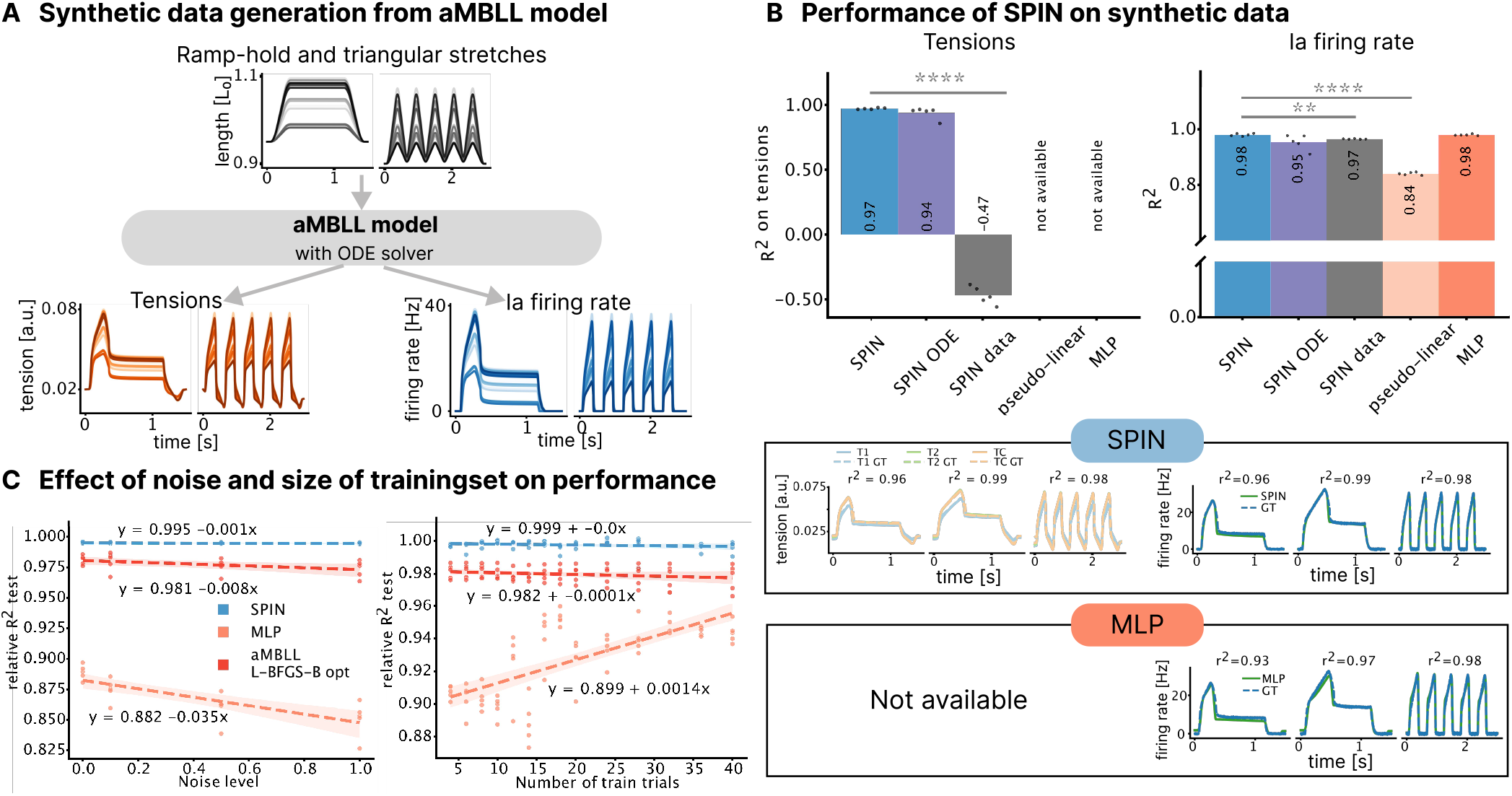
Proof of concept of SPIN training using synthetic data. **A** Synthetic data were generated by solving the differential equations of the aMBLL model, providing reference tension and firing rate profiles across a range of muscle stretches. **B** We trained the SPIN model, along with baseline comparison models, on the synthetic data with additive Gaussian noise (0.02). The *R*^2^ scores for tension and firing rate predictions were computed across five folds (top panels), with the mean *R*^2^ values indicated on each bar. Statistical significance relative to SPIN was determined using a paired two-sided t-test. Predicted tension and firing rates for three example test trials are shown, demonstrating that SPIN accurately predicts both tension (top left) and firing rate (top right). In contrast, the MLP model fits the firing rate (bottom right) but does not predict tension. *R*^2^ scores for the shown trials are also provided. **C** Performance of SPIN, MLP model and aMBLL model optimzied with L-BFGS-B oprimizer when trained with varying number of trials and noise levels. The *R*^2^ scores (relative to the maximum *R*^2^ achieved on the test set) were computed for models trained with varying noise levels (left) and different numbers of training trials (right), using the same test set (20 trials without noise). Models were trained on five folds, with each dot representing a fold, and the linear regression fit shown for both SPIN and MLP. For the noise comparison (left panel), five training trials were used, and for the trial count comparison (right panel), a noise level of 0.02 was added to each trial.

We evaluated the goodness-of-fit of the model using the *R*^2^ metric for both the firing rate and the tensions of the three intrafusal fibers on the test dataset (Figure 2B). The baseline MLP model achieved high accuracy (*R*^2^ = 0.98; N=5 folds) in fitting the firing rate but lacked structural fidelity (for example, it has no predictions of the intrafusal fiber tensions). The pseudo-linear model, with a lower complexity than the MLP, showed a reduced fit (*R*^2^ = 0.84; N=5). For the SPIN model, we evaluated three variations: the full model trained with the total loss, a model trained exclusively on the ODE loss, and one trained only on the data loss (Eq. (4)). A combination of both losses was essential to achieve accurate fits for both the firing rate (*R*^2^ = 0.98; N=5) and fiber tensions (*R*^2^ = 0.97; N=5).

Notably, the SPIN model trained solely on the data loss fit the firing rate well (*R*^2^ = 0.97; N=5) but failed to accurately predict fiber tensions (*R*^2^ = −0.47; N=5). This underscores the essential role of the ODE loss in constraining the tensions produced by the neural network. Without the ODE loss, the model can fit the firing rate data without maintaining the biophysical accuracy of the fiber dynamics. By incorporating the ODE loss, SPIN ensures that the predicted fiber tensions align with known physiological dynamics, resulting in a model that not only replicates the predictions of a mechanistic model but also captures the underlying muscle spindle behavior in a more comprehensive and biologically faithful manner.

We assessed the impact of (simulated) noise in the training data and the number of training trials on model performance (Figure 2C), comparing SPIN to a data-driven model (MLP) and a mechanistic model utilizing an ODE solver with parameter optimization (aMBLL with L-BFGS-B). We compute the *R*^2^ score on the same test set (without noise) for the models trained with varying noise levels and number of training trials. A key advantage of the SPIN model is its ability to be pre-trained using only ODE collocation points, without relying on measured firing rate data. As a result, even with a small number of training trials and high noise levels, SPIN was able to predict the firing rate with high accuracy (relative *R*^2^ = 0.99). In contrast, the MLP model was more sensitive to both noise and the number of training trials. Its performance declined with increasing noise (slope =−0.035; *p* = 0.598) and improved with additional training trials (slope = 0.0014; *p* = 0.462). In comparison, SPIN showed minimal sensitivity to these factors, with a near-zero slope for noise (−0.001; *p* = 0.147) and a small positive effect for the number of trials (slope = 0.0; *p* = 0.032) for the number of trials. The aMBLL model with parameter optimization also showed low sensitivity to noise and trial count, while achieving higher performance than the MLP. Indeed, the aMBLL model inherently incorporates mechanistic properties and only requires parameter tuning to fit the new data.

These experiments demonstrate that the SPIN model can be trained with noisy and sparse data to outperform traditional data-driven models of muscle spindles, while maintaining the mechanistic components of the aMBLL model. This training process is faster than directly optimizing the parameters in the aMBLL model. Moreover, compared to purely data-driven machine learning models, SPIN, as a physics-informed neural network, requires less training data. The physical constraints embedded in the model guide it toward an accurate solution, even with limited observations, improving its robustness and generalizability. This is particularly important for muscle spindles, as it is often not possible to maintain a spindle afferent recording for a long time during naturalistic movement.

### SPIN allows for cross-species comparisons

We successfully trained the SPIN model using synthetic noisy data to replicate firing rate and tension predictions from the baseline mechanistic model. We next evaluated SPIN on a range of experimental data.

With the development of microneurography studies targeting human muscle spindles, important questions on the variability of muscle spindle function across mammals have been raised (22). Notoriously, human muscle spindle afferents fire at a considerably lower rate than cat afferents (22). It remains unknown whether the mechanistic models developed based on observations in cat muscle spindles are still applicable to human muscle spindles. It is not clear whether the only difference is in the parameters of the models or if there are more fundamental variations that cannot be captured by the existing models. For this purpose, we examined the predictions of SPIN on Ia afferent responses during passive stretches of the *triceps surae* muscle in anesthesized cats (55) and rats (35) and *extensor capri radialis* Ia afferent responses during active wrist movements in humans (58).

We trained the SPIN model on Ia afferent responses, utilizing data from multiple Ia afferents (11 from cat data (one having insufficient data), 10 from rat data, and 8 from human data). Each SPIN model was trained on individual afferents, allowing the mechanotransduction parameters *λ*_*m*_ and three of the fiber parameters *λ*_*f*_ to vary. As in numerous previous studies (34, 56), we estimated the normalized fascicle length (the relevant input to the aMBLL and SPIN models) from the experimentally recorded muscle-tendon lengths in cats and from joint angles and a musculoskeletal model in humans (see Methods). We compared the performance of SPIN against the SPIN model trained solely with data loss, the pseudo-linear model and the aMBLL model with parameters optimized with L-BFGS-B (Figure 3).

**Figure 3.**
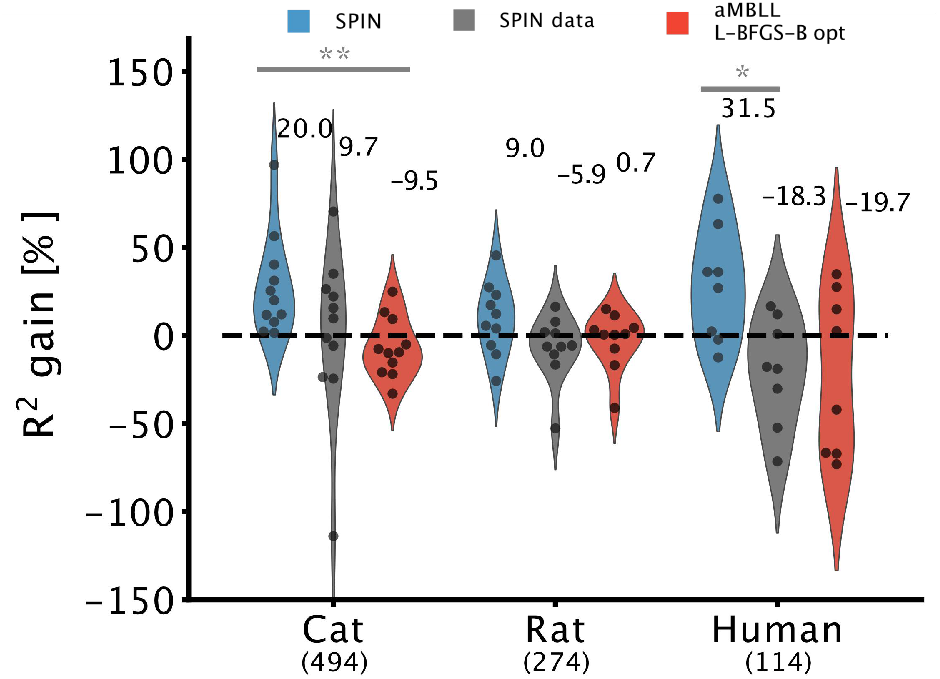
Distributions of *R*^2^ gain (in %) with respect to the baseline pseudo-linear values. We plot the gain for the three datasets; each point corresponds to an afferent (mean over the folds). We compare *R*^2^ gain between the SPIN model (blue), SPIN data (grey) and the optimized aMBLL model (red). Under each label we indicate the total number of test trials considered for each dataset. Statistical significance with respect to SPIN were calculated using paired two-sided t-test. We display the median value for each model.

First, we compared the overall performance of SPIN, SPIN data, and the optimized aMBLL across all afferents (Figure 3) focusing on the distribution of the gain in *R*^2^ for each trained model relative to the pseudo-linear model. We considered the pseudo-linear model as baseline, as it is similar to previous models (26, 27, 29, 30, 54, 55). The (ODE-fitted) aMBLL model showed positive *R*^2^ gains in only 11 out of 29 afferents, indicating that in most cases, the pseudolinear model—a purely statistical approach—fit the data better. This highlights the challenges in effectively optimizing the ODE-based aMBLL model. In contrast, SPIN outperformed the pseudo-linear model for all afferents except for 4/10 rat and 2/8 human afferents, demonstrating SPIN’s superior representational capacity over the statistical pseudolinear model and its stronger training capabilities compared to the optimized aMBLL model. Across species, SPIN consistently outperformed the optimized aMBLL model in terms of accuracy. For the cat data, SPIN showed a median performance improvement of 20.0%, compared to −9.5% for aMBLL, this difference being statistically significant. In the rat data, SPIN also had a higher median gain of 9.0%, compared to 0.7% for aMBLL. The difference in human data was more pronounced, with SPIN achieving a median improvement of 31.5% versus −19.7% for aMBLL.

Next we looked at the afferent-specific fits. All models trained on the cat and rat data generally produced strong fits (up to *R*^2^ = 0.9). In contrast, the fit for human data was lower, with *R*^2^ values ranging from 0.2 to 0.6, likely due to differences in experimental conditions. The cat and rat data, recorded from anesthetized animals with muscle stretches generated by a servo motor, exhibiting consistent stretches, leading to stereotypical responses: a large increase at stretch, slight decay during hold, and a large drop at release (Figure 4B). These consistent responses are easier to model. In contrast, the human data, recorded during active movements, involved more naturalistic muscle stretches and exhibited greater variability in responses (Figure 4B right panel). Furthermore, there is evidence that during active voluntary movements, the spindle’s sensitivity to length changes is reduced (22, 59). In particular, during shortening, most of the muscle spindles are silenced (as evidenced by the bottom panel of Figure 4B on the right); it is the spindles in the lengthening antagonist muscle which encode the muscle stretch information (60). Given that the spindle recordings in human active movements are from a single muscle, this could make fitting the firing rate more difficult.

**Figure 4.**
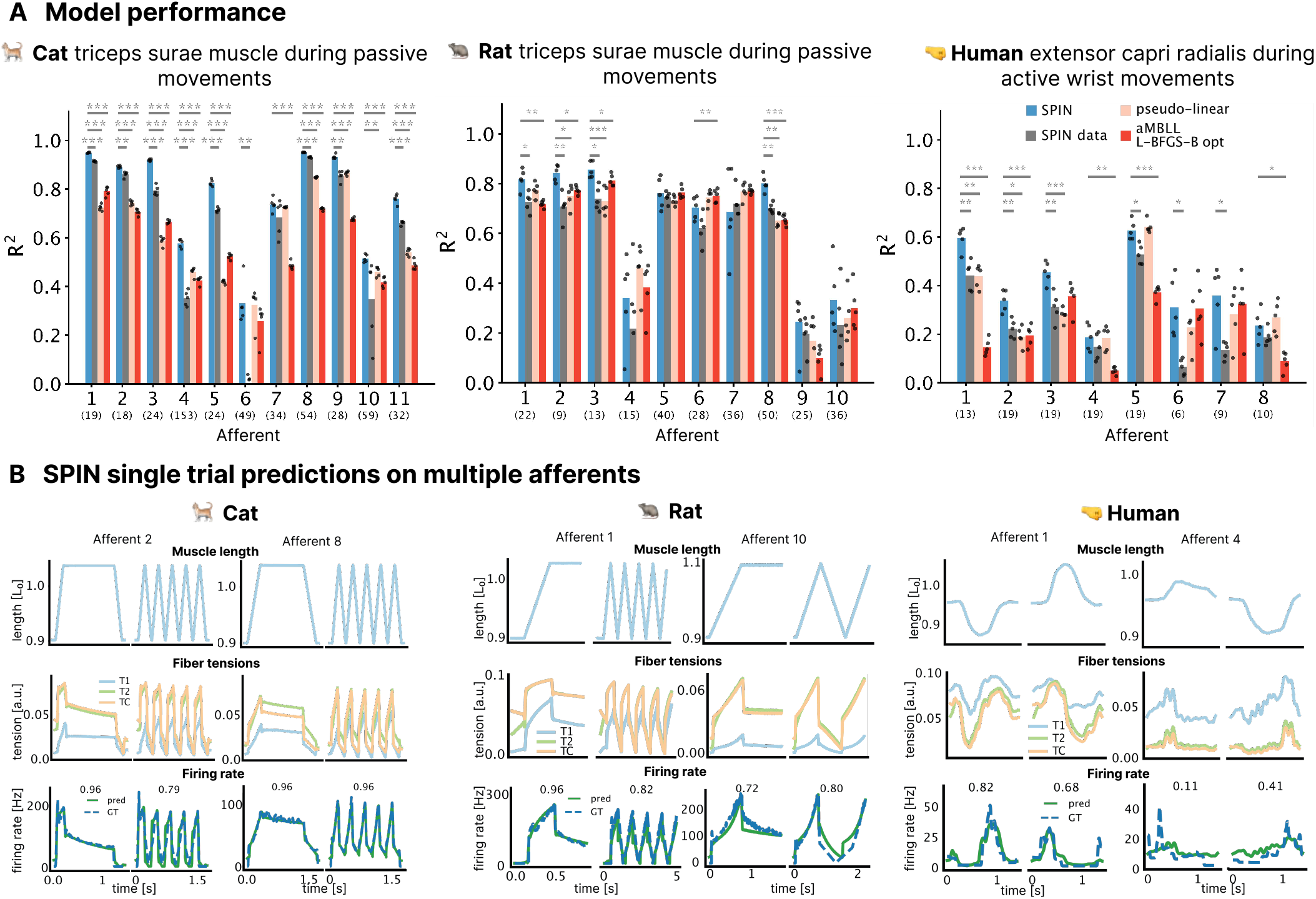
SPIN can effectively be tuned to experimental data. **A** Performance of SPIN compared to 3 baseline models trained on three datasets: cat triceps surea muscle during passive movements (left), rat triceps surea muscle during passive movements (middle), human *extensor capri radialis* muscle during active wrist movements (right). We train a model on each individual afferent and compute the *R*^2^ coefficient on test data for 5 different folds. Each point corresponds to a different training fold. The bar represents the mean value. We compare SPIN (in blue) to SPIN data, model trained with no ODE loss (grey), pseudo-linear (orange), aMBLL optimized with L-BFGS-B (red). The numbers below each afferent correspond to the number of trials in the test set. Statistical significance with respect to SPIN was calculated using paired two-sided t-test. **B** Single trial predictions from SPIN model on multiple afferents. For each dataset cat (left), rat (middle), human (right), we plot the fascicle length (top), predicted fiber tensions (middle) and firing rate prediction (green solid) and ground truth (blue dashed) (bottom). Above the firing rate predictions, the *R*^2^ coefficient for the trial is indicated. The different afferents represented show the diversity of afferent response even for similar muscle stretches.

SPIN outperformed both baseline models in all but two cat afferents (6 and 7). In rat and human data, SPIN showed greater variability in performance, with a larger range in *R*^2^ scores and less consistent improvement over the baseline models (Figure 4A).

Given that the aMBLL model and baseline parameters were based on the anatomy and function of cat muscle spindles, it is perhaps unsurprising that SPIN best fits cat data. The afferent data from cats and rats were recorded under similar conditions, though the muscle stretch profiles for rats contained more complex trials. The *R*^2^ values of all models trained on cat afferents tend to exhibit a smaller variance than those trained on rat afferents. Since we initialize the SPIN model parameters with the baseline models provided by Mileusnic et al. (34) for cats, SPIN might struggle to find appropriate parameters for the rat afferents. Additionally, there may be fundamental differences between the species that cannot be captured by the current mechanistic model alone.

The updated version will compare parameters across species and highlight how SPIN can model fusimotor inputs (*γ*), stay tuned.

## Discussion

Prior models of muscle spindles lacked either physical fidelity or computational tractability with some relying on transfer functions that neglect underlying physical mechanisms, while others, though more structurally realistic, lack computational efficiency and scalability. We presented a model of muscle spindles that combines structural fidelity with computational efficiency, leveraging the power of physics-informed neural networks. By integrating principles of biomechanics and neural dynamics, our model captures the interplay of biomechanics and mechanotransduction processes within muscle spindles while maintaining computational tractability.

Importantly, adding a differential equation loss has two implications. It introduces interpretable variables in otherwise black-box neural network models and it constrains the behavior of those variables. It is important to realize that, *per-se*, makes the loss harder to minimize. It will only be easier to satisfy the loss if the equations are (approximately) correct. We have seen evidence for this in this study when fitting SPIN across species.

Overall, we found that SPIN captures muscle spindle data better than previous models, while also being fast and obeying mechanical and mechanotransduction equations. Proprioception is essential for planning and executing precise motor actions and relies on sensory signals from mechanosensory neurons distributed within muscles, tendons, and joints. Muscle spindles convey information through sensory afferents to the central nervous system. While traditionally viewed as stretch receptors encoding muscle length and velocity, recent insights suggest they may have a role as signalprocessing devices, adapting responses to task demands via spinal gamma motor neurons (24). Accurate and scalable models like SPIN are key to further gain insights into these hypotheses. First, these more precise muscle spindle models can be incorporated into both normative and data-driven models of the proprioceptive pathway (9, 61, 62), potentially enhancing explanatory power beyond the spinal cord. Additionally, integrating these models into frameworks for motor control and learning (63, 64) could offer valuable insights into how sensory feedback informs movement and adaptation.

### Natural extensions

Here we considered one specific ODE in SPIN, namely we built on the the classic aMBLL equation. In future work, one could explore alternative differential equations and ideally in a back and forth with experiments improve our understanding of muscle spindles.

Firstly, the aMBLL model is structurally non-identifiable, meaning that we can achieve the same fit to the firing rate data with different set of parameters (65). One could use SPIN variants to propose experimental stimuli to discern different parameters in a high-throughput fashion.

Secondly, the simple mechanotransduction module makes it hard to model history effects and certain dynamics. Namely, in the cat and rat data one can observe history dependent response with initial very large peak (after rest) (37, 66). This cannot be explained by the aMBLL ODE. The firing rate decay during the hold phases can be captured up to a certain point but could be improved if we modeled some dynamics of the channel openings (67). Recent studies have explored the influence of cross-bridge dynamics on spindle responses (35, 37). Indeed, we plan to expand the mechanotransduction equations in SPIN.

Thirdly, Blum et al. (55) argued that primary muscle spindles preferentially encode muscle force-related variables vs. stretch-related variables. Concretely, they showed that recorded afferents in cat *triceps surae*, in response to imposed passive stretches, could be better predicted by a pseudo-linear model based on muscle force and yank (rate of change of force) rather than by a model based solely on muscle kinematics. One could also input muscle force and yank into SPIN and we expect better predictions. However, a more interesting extension would be to integrate the force components in the ODE.

Another interesting direction is to combine our approach with automatic methods for ODE discovery, such as sparse regression (68, 69), or symbolic regression (70, 71).

Interpretable methods are of great interest, as they facilitate multiscale modeling (72), for example linking data from molecular levels, such as transcriptomics, to higher-level processes like electrophysiology (73). While PINNs have gained significant traction in physics (38), their application in biology has been relatively limited (65, 74–76). With SPIN, we demonstrate the potential for developing interpretable, statistically robust models in computational neuroscience. Beyond modeling muscle spindles, we believe that the approach used in SPIN could be applied across different scales of neural dynamics.

## Methods

### The adapted Mileusnic-Brown-Lan-Loeb model

We constructed a baseline mechanistic spindle model to establish the physical constraints for the SPIN model. This model, called aMBLL, is adapted from the MBLL model proposed by Mileusnic et al. (34) and the Hasan model (31).

The main component of these models are the tension equation, describing the dynamics of the tension of the intrafusal fiber tensions. The main difference between the two models is in the definition of the parameters, the units, the normalization of the muscle kinematics and the addition of the mass term in the MBLL model.

Adding the mass term in the MBLL model results in a second-order ODE. In a more recent paper, Chacon et al. (77) find that the MBLL model without mass results in the same prediction performance and shows more robustness of the solution (77). Hence, we simplify the MBLL equation to the mass-free case.

The tension in the sensory zone is given by:

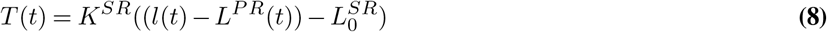

and the tension in the polar zone is

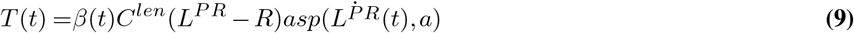

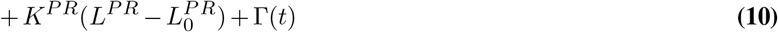

where *asp*(*x, a*) = *sign*(*x*) |*x*| ^*a*^. Given that the two tensions are equal we can use the first equation to solve for *L*^*P R*^(*t*) and replace it in the second equation. We get

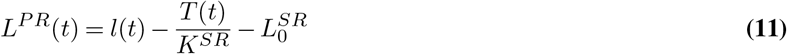

and the following differential equation for the tension

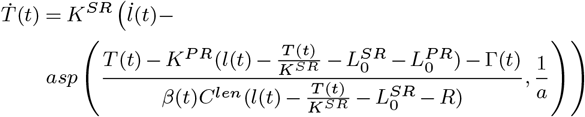

Defining 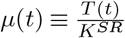 we can write the tension equation as

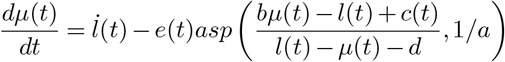

with

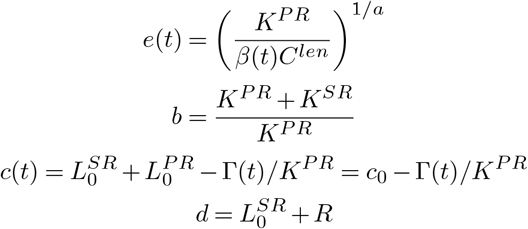

This form is very similar to the expression in Hasan (31):

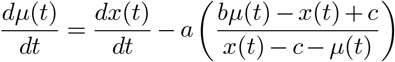

The advantage of the MBLL model is that the effect of *γ* fusimotor inputs is explicit in the model (*β*(*t*) and *τ* (*t*)). From now on we use this adapted, mass-free MBLL model (aMBLL). We can write the equation as

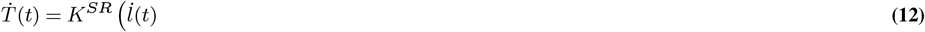

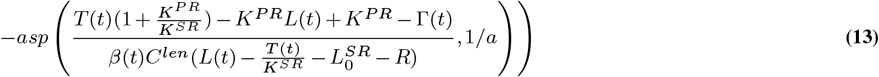

where we have used the fact that 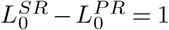. *β*(*t*) and Γ(*t*) depend on the fusimotor inputs as follows

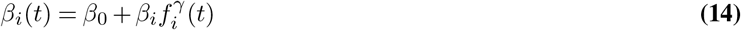

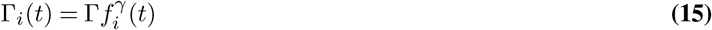

where 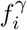 has values between 0 and 1 representing the activation from the *γ* fusimotor input. Bag 1 only receives inputs from dynamic *γ* and bag 2 and chain only receive static *γ* inputs.

We can define the right hand side of the above equation as

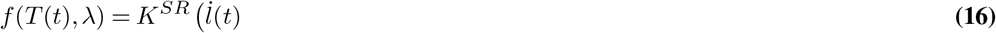

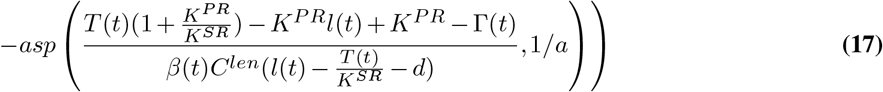

where *λ* denotes the 11 parameters in the equation.

Overall, the aMBLL model takes five inputs: fascicle length (*l*), velocity (*v*), acceleration (*a*), dynamic *γ* activation (*γ*_*d*_), and static *γ* activation (*γ*_*s*_), and outputs the firing rates of Ia and II afferents. It includes three types of intrafusal fibers—bag1, bag2, and chain—each governed by the tension differential equation:

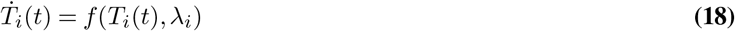

where *T*_*i*_ represents the tension of fiber *i, λ*_*i*_ denotes the fiber’s parameters, and *f* is defined as in Eq. (17). Although the tension equation is consistent across all fiber types, the parameters *λ*_*i*_ vary:

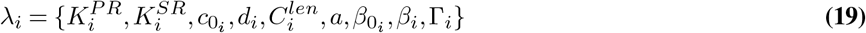

The mechanotransduction module uses these fiber tensions to compute the afferent firing rates as defined in Eq. (2) and Eq. (3). The firing rate output depends on the following mechanotransduction parameters:

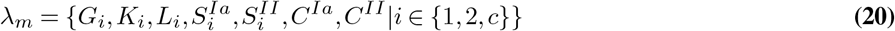

To simulate the aMBLL model, we solved the tension equation for each fiber type using the `scipy.integrate.odeint` function with absolute error tolerance *atol* = 10^−8^ and relative error tolerance *rtol* = 10^−6^. Unless otherwise specified, initial tension values were set to *T*_*i*_(0) = 0.02.

We used the reference parameter values from the cat soleus data in Mileusnic et al. (34) summarized in table 1.

**Table 1.** Summary of reference values for the aMBLL model parameters and the bounds used for the optimization.

| Param name | bag 1 | bag 2 | chain | Bounds |
| --- | --- | --- | --- | --- |
| $K^{PR}$ | 0.15 | 0.15 | 0.15 | [0.1, 0.2] |
| $K^{SR}$ | 10.4649 | 10.4649 | 10.4649 | [1, 20] |
| $a$ | 0.333 | 0.333 | 0.333 | [0.1, 0.9] |
| $c_0$ | 0.8 | 0.8 | 0.8 | [0.6, 1] |
| $d$ | 0.5 | 0.5 | 0.5 | [0.4, 0.6] |
| $C^{len}$ | 1, 0.42 | 1, 0.42 | 1, 0.42 | [0.1, 1.0] |
| $\beta_0$ | 0.0605 | 0.0822 | 0.0822 | [0.001, 0.2] |
| $\beta$ | 0.2592 | -0.046 | -0.069 | - |
| $\Gamma$ | 0.0289 | 0.0636 | 0.0954 | [0.01, 0.5] |
| $L_m$ | 0.0023 | 0.0023 | 0.0023 | [0, 0.05] |
| $K_m$ | 10.46 | 10.46 | 10.46 | [5, 15] |
| $G$ | 20000 | 10000 | 1000 | [5000, 300000] |
| $S^{Ia}$ | 0.2 | 0.2 | 0.2 | [0, 1] |
| $C^{Ia}$ | 0 | - | - | [0, 50] |

### SPIN for modeling muscle spindles

SPIN employs a feed-forward neural network (NN) to model the dynamics of intrafusal fibers. Following a hyperparameter search, we identified an optimal architecture consisting of three hidden layers, each with 128 units. The NN receives six inputs: fascicle length (*l*), velocity (*v*), acceleration (*a*), dynamic *γ* activation (*γ*_*d*_), static *γ* activation (*γ*_*s*_), and time (*t*), and produces three outputs corresponding to the tensions of the bag1 (*T*_1_), bag2 (*T*_2_), and chain (*T*_*c*_) fibers.

The NN uses a hyperbolic tangent (tanh) activation function for all layers except the output layer, which employs a sigmoid activation function. Weights are initialized using Xavier initialization, while biases are set to zero. The fiber parameters (*λ*_*f*_) and mechanotransduction parameters (*λ*_*m*_) were initialized to the reference values listed in Table 1, ensuring the model begins from a biologically plausible configuration.

Each input to the NN is scaled based on predefined bounds derived from experimental data (Table 2). To improve training stability, the NN outputs are scaled by 0.1, reflecting the approximate tension range of 0 to 0.1 in the model.

**Table 2.** Upper and lower bounds on the inputs to the NN.

| Input | $l$ | $v$ | $a$ | $\gamma_d$ | $\gamma_s$ | $t$ |
| --- | --- | --- | --- | --- | --- | --- |
| Lower bound | 0.8 | -1 | -50 | 0 | 0 | 0 |
| Upper bound | 1.3 | 1 | 50 | 100 | 100 | 4 |

### SPIN Model training

The model training process involved optimizing the weights of the neural network (NN) along with parameters governing the fiber and mechanotransduction dynamics. We train a model for each afferent, obtaining the parameters of the model that fit the specific afferent best.

When training on experimental data, we initialized the NN weights to the value found when training on synthetic data. This effectively acts as a pre-training of the NN weights such that the tension outputs of the NN satisfy the ODE equations with the baseline model parameters. Similar performance could be achieved training from weights initialized with Xavier but more hyperparameter optimization was required and the variance in the results was greater.

Training was performed with a batch size of 1, corresponding to a single experimental trial. For each batch, the total loss was computed by combining the data loss, the ODE loss, and the initial conditions loss as defined in Eq. (4). The data loss ℒ_*d*_ was computed over all the time points of the trial, with *N* ^*d*^ = *T*_*trial*_*/dt* where *T*_*trial*_ is the duration of the trial in seconds and *dt* is the timestep in seconds. For the ODE loss calculation ℒ_*ODE*_, we generated *N* ^*o*^ = 10, 000 collocation points, *x*_*n*_, by sampling for each input a value from a uniform random distribution within the bounds defined in the previous subsection. This approach ensured that the model not only accurately fit the observed data but also generalized well to unseen conditions within the physiological range. The weight *w*_*ODE*_, which determines the contribution of the ODE loss relative to the data loss, was identified as a critical hyperparameter and was tuned using grid search. Details on the grid search are specified below.

We trained the SPIN model using the Adam optimizer on the neural network weights *θ* and the model parameters (fiber *λ*_*f*_ and mechanotransduction *λ*_*m*_). In some cases, different learning rates were assigned to the NN weights and the model parameters to account for their varying scales and sensitivities during optimization. These learning rates *α*_*NN*_, *α*_*f*_ and *α*_*m*_ were treated as hyperparameters and optimized by grid search (Table 3 summarizes the values used). A learning rate scheduler was used to reduce the learning rates by 10% every 5 epochs.

**Table 3.** Summary of hyperparameters used for SPIN models trained. Model refers to the model type as it appears in the figures. Data refers to the type of data used for training. Weight init indicates if we used the default Xavier initialization or the weights from the trained model on synthetic data.

| Model | Data | Weight init | Learning rates | $w_{ODE}$ | $w_{IC}$ | L-BFGS |
| --- | --- | --- | --- | --- | --- | --- |
| SPIN | Synthetic | Xavier | [0.0001, 0.0001, 0.0001] | 10 | 1 | Yes |
| SPIN ODE | Synthetic | Xavier | [0.0001, 0.0001, 0.0001] | 10 | 1 | Yes |
| SPIN data | Synthetic | Xavier | [0.0001, 0.0001, 0.0001] | - | 1 | NO |
| SPIN pre-trained | Synthetic | from pre-trained | [0.0001, 0.001, 0.01] | 10 | 1 | NO |
| SPIN | Experimental | from pre-trained | [0.0001, 0.001, 0.01] | 1 | 10 | NO |
| SPIN data | Experimental | Xavier | [0.0001, 0.0001, 0.0001] | - | 1 | NO |

At each training step, we applied gradient clipping (clipping to a maximum of 1) and restricted certain parameters: *C*^*Ia*^ and *L*_*i*_ were constrained to be positive, while *S*_*i*_ was limited to the range of 0 to 1.

For synthetic data, when training the models SPIN and SPIN ODE, we also applied an L-BFGS optimizer every 5 epochs of the Adam optimizer. We found that the L-BFGS optimizer did not improve performance training significantly when using a pre-trained SPIN model or when training the SPIN data model.

The model was trained on the training set and performance was monitored using the validation set. To prevent overfitting, we implemented an early stopping criterion based on the validation loss. The training procedure included adjustment of the learning rate and early stopping. Every 5 epochs where the validation loss did not decrease, the learning rate was reduced by 20%. Training was terminated if the learning rate was decreased 100 times without improvement in validation loss. We trained over a total of 800 epochs, although early stopping was often reached.

### Training baseline models

The baseline models were trained to minimize the *L*_2_ loss between the predicted firing rate and the ground truth firing rate. We computed the mean square error per trial except for the pseudo-linear model where we could stich all train trials together.

#### aMBLL

We trained the aMBLL model by optimizing its parameters using the L-BFGS optimization algorithm. Two scenarios were considered: (1) when only the mechanotransduction parameters (*λ*_*m*_) were optimized, and (2) when at least one fiber parameter was included in the optimization.

In the first scenario, the model was simulated by solving the ODE for each trial *i* in the training dataset, saving the predicted tensions (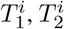 and 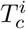). These pre-computed tensions were used in the optimization process, allowing for the predicted firing rate to be computed using equations Eq. (3) and Eq. (2). This approach significantly improved computational efficiency, as the ODE system did not need to be solved during each iteration of the optimizer, only once before the optimization. In the second case, the ODE equations need to be solved at each iteration since the tensions depend on the fiber parameters that are optimized.

A sensitivity analysis was performed to identify the parameters that most strongly influence the dynamic behavior of the aMBLL model. Unless otherwise specified, we optimized the following parameters: *S*_1_, *S*_2_, *C, K*_1_, *K*_2_, *G*_1_, and *G*_2_. Although we attempted to optimize simultaneously mechanotransduction and fiber parameters (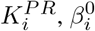, and 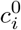), this resulted in excessive computation times with minimal performance improvement. Thus, we resolved to only optimization of the mechanotransduction parameters.

As reported by Mileusnic et al. (34), the bag2 and chain fibers contribute equally to the overall firing rate in the absence of gamma fusimotor input. Based on this finding, we assumed that the fiber tensions and parameters were equal in these cases, which reduced the number of parameters to optimize and simplified the overall process.

For each training, we performed five separate optimizations with different initial parameter values, drawn from a uniform random distribution within the bounds specified in Table 1. The parameter set that yielded the lowest error was selected for the final model.

#### Pseudo-linear

The pseudo-linear model is a linear model with a half-wave rectifier activation function to capture non-linearities in the input-output relationship. The model outputs a weighted sum of the rectified inputs with an additional intercept term:

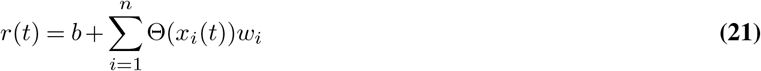

where *b* is the intercept, *n* is the number of inputs *w*_*i*_ the weight for each input channel *x*_*i*_ and Θ(*x*) = *max*(0, *x*) is the half-wave rectifier function.

The pseudo-linear model was trained to output the firing rate given three inputs: fascicle length (*l*), velocity (*v*), acceleration (*a*). We optimized the weights *w*_*i*_ and the intercept *b* with the function `scipy.optimize.minimize` using the L-BFGS-B algorithm. The weights were initialized to 1 and the bias to 0.

#### MLP

We implemented a multi-layer perceptron (MLP) with an architecture identical to the neural network used in the SPIN model. The MLP consists of three hidden layers, each containing 128 units, and uses the hyperbolic tangent (tanh) activation function for all hidden layers. The MLP takes six inputs: fascicle length (*l*), velocity (*v*), acceleration (*a*), dynamic *γ* activation (*γ*_*d*_), static *γ* activation (*γ*_*s*_), and time (*t*) and outputs the predicted afferent firing rate.

Each input was normalized according to predefined bounds derived from experimental data to ensure consistency with training for SPIN (Table 2). The output layer used a linear activation function and was scaled by a factor of 10 to improve convergence during training.

The model weights were initialized using Xavier initialization to ensure proper scaling of inputs through the network layers, while the biases were initialized to zero. Training was conducted using the Adam optimizer with a learning rate *α*_*MLP*_ optimized through grid search. A learning rate scheduler was applied, reducing the learning rate by 10% every 100 epochs to improve stability and convergence.

The training set is divided in batches of size 16 except for the simulations presented in Figure 2C where a smaller batch size of 2 was used due to the limited number of training trials. To prevent overfitting, we employed early stopping, using a validation set identical to that used in the SPIN model training process. The model was trained for a maximum of 5000 epochs, although most simulations achieved early stopping, halting the training when the validation loss ceased to improve.

### Hyperparameter optimization

We performed hyperparameter optimizations on the SPIN and MLP models with `wandb`. We ran 2 cross-validation folds for each hyperparameter optimization and used the validation loss (averaged over the last 10 epochs) as the metric.

We did a first round of hyperparameter search to find general hyperparmeters for SPIN to work on synthetic and experimental data. Using a grid search with 3 folds on the following hyperparameters: number of layers, number of units in each hidden layer, learning rate of the NN weights, number of epochs, *w*_*ODE*_, *N* ^*o*^, and whether to use the LBFG-S optimizer or not.

In order to compare accurately the performance of SPIN and and baseline models on synthetic data, we performed a second round of hyperparameter optimization. We performed Bayesian hyperparameter optimization on synthetic data with added noise with *σ* = 0.02 and 5 training trials. The optimization was done on the final validation loss (averaged over the last 10 epochs).

For the SPIN model, we optimized the following parameters: **N**^**o**^, the number of collocation points for the ODE loss, the number of times we decrease the learning rate before stopping training, the learning rate *α*_*θ*_ for the NN weights, the use of LFGS optimizer or not, the percentage decrease of learning rate in the validation loop, and *w*_*ODE*_, the coefficient weighting the proportion of ODE loss.

For the MLP baseline model, we optimized the following parameters: the size of the training batch, the learning rate, the percentage decrease of learning rate in the validation loop, and the number of times we decrease the learning rate before stopping training.

We use the optimal hyperparameters to train the respective models with varying noise and number of train trials.

### Cross-validation

Unless otherwise specified, we performed a 5-fold cross-validation to evaluate the performance of the models across different subsets of the data. The dataset was split into training, validation, and test sets with default proportions of 0.2 for the test set and 0.1 for the validation set (relative to the test/train split). This process ensured that each fold had an independent test set, while the validation and training sets were used for model optimization.

We split the data ensuring that all experimental conditions were represented in both the training and test sets. For that purpose, we had to adjust the proportions of the test and validation sets for datasets where certain conditions were underrepresented, requiring an increase in the fraction of data allocated to the test and validation sets. Specifically, for the cat and rat dataset, we set the proportion of test data to 0.4 and of validation set to 0.2.

The model’s performance was evaluated on the test set for each fold using the *R*^2^ coefficient as the primary metric. The *R*^2^ was calculated as follows.(1) The test data across all conditions and trials were flattened into a single vector. (2) any missing values present in either the predicted or ground truth firing rate data were removed. (3) The score was calculated on all points in the flattened test data vector using the function `sklearn.metrics.r2_score` between the two vectors. This approach provided a single performance metric for each fold and ensured that the value reflected the model’s overall ability to predict the test data, regardless of the specific trial length.

To compare the performance of various models on experimental data, we computed the gain in test set *R*^2^ values relative to a baseline model (pseudo-linear model):

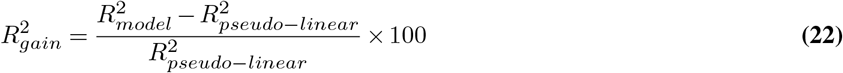

This gain was calculated for each afferent, fold, and model type across different datasets. For each type of model, each dot represents the value for each fold, for each afferent. Paired two-sided t-tests were performed between the results for the SPIN model and the rest of the models.

### Quantification and statistical analysis

Statistical significance was evaluated with paired two-sided t-tests using the function `scipy.stats.ttest_rel`. Significance levels were annotated on the plot using asterisks to indicate p-value threshold. Not significant (ns) for *p >* 0.01; * for *p <* 0.05; ** for *p <* 0.01; *** for *p <* 0.001, unless specified otherwise.

### Generating synthetic data

We generate synthetic data for the model identification analysis of SPIN. We first generate sample muscle stretches producing from which we compute the fascicle length, velocity and acceleration.

Using these inputs and default *γ*_*d*_ = *γ*_*s*_ = 0, we generate the ground truth firing rates by simulating the aMBLL model. For each trial, we solve the fiber ODE equation to obtain each tension and then compute the Ia and II firing rate. We simulate the aMBLL model as described in the section “The adapted Mileusnic-Brown-Lan-Loeb model”.

We generate 20 trials for each of 5 types of muscle stretches *l*(*t*):

1. Ramp-hold-release stretches of length 1.5s. The trials start at constant fascicle length 0.95, initializing a stretch of length 0.2s at 0.1s and initializing the release at 1.2s for 0.2s back to length 0.95. The length of the hold phase is randomly chosen between 0.98 and 1.1.
2. Ramp-hold-release as (1) except varying length of stretch between 0.15s and 0.4s and constant hold value of 1.08.
3. Ramp-hold-release stretches of length 1.5s. All as (1) except varying length of release between 0.1s and 0.4s and constant hold value of 1.08
4. Ramp-hold-release as (1) except varying starting fascicle length between 0.9 and 1.0 and constant hold value of 1.08.
5. Triangular stretches with duration 3s. The initial value is 0.9, there are 5 peak times with varying peak value between 0.95 and 1.1.

The timestep for each trial is *dt* = 0.0001*s*. We filter the fascicle length with a savgol filter with window size 0.1*/dt* + 1, with polynomial order 1. We compute the velocity and acceleration with a savgol filter on the unfiltered fascicle length with the window size 0.1*/dt* + 1 and polynomial order 2 for the velocity and 3 for the acceleration.

We add noise to the firing rate of the training trials. For each trial, noise is drawn from a random Gaussian distribution with mean 0 and standard deviation equal to the product of the noise level and the power of the firing rate.

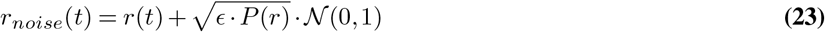

where *r*(*t*) is the firing rate, *ϵ* is the noise level, 𝒩 (0, 1) is the standard random normal distribution and 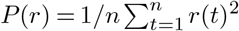 is the power of the signal, *n* is the number of timepoints in the trial. This procedure is applied independently to each trial to simulate variability in neural firing patterns.

### Afferent Firing Rate Datasets

The afferent firing rate data from different sources were processed for use in our model.

#### Cat Data

We obtained cat afferent data from Blum et al. (78). The firing rate was calculated from recorded spike times using inter-spike intervals, and subsequently interpolated over the entire trial duration using a linear filter. Fascicle length was estimated as described in the “Estimating the Muscle Fascicle Stretch from Whole Muscle Stretch” section below. Fascicle velocity was calculated by scaling the recorded velocity according to the fascicle length coefficient. Fascicle acceleration was derived from the filtered velocity. We applied a low-pass filter with a cut-off frequency of *f*_*c*_ = 40Hz to both velocity and acceleration signals. All signals had a 2kHz sampling frequency.

The “condition” of each trial corresponds to the trial stretch type indicated in the dataset. There were 5 different conditions: accel-series-proc, trad-ramp-proc, trad-series-proc, accel-ramp-proc, triangle-series-proc. The first 4, were ramp-hold-stretch profiles with different speeds and the last one had triangular stretch profiles. Of the 12 recorded afferents, we included 11, excluding one afferent due to having fewer than 40 recorded trials.

#### Rat Data

We obtained rat afferent data from Blum et al. (79). The firing rate was provided as the instantaneous firing rate, which we interpolated to ensure a continuous value at each timestep.

Due to variations in sampling frequencies between experiments (e.g., 2kHz for animals ‘0’ and ‘1’, and 10kHz for animals ‘4’ and ‘5’), we downsampled all data to a uniform 2kHz frequency. Fascicle length was estimated as described in the “Estimating the Muscle Fascicle Stretch from Whole Muscle Stretch” section. Velocity was computed from the recorded values, scaled by the same factor as muscle length, and filtered with a low-pass filter (cutoff 40Hz). Acceleration was computed from the filtered velocity and filtered with a low-pass filter (cutoff 40Hz).

The dataset included 9 conditions: accel-ramp, triangle-series, accel-series, trad-series, ramp, amptri, timetri, pseudorand, trad-ramp. To maintain consistency in data splitting (train/validation/test), we excluded conditions (trad-ramp and accel-series) with only a single trial for any afferent, resulting in the removal of four trials. Additionally, trials exhibiting missing spike times during the latter half were truncated at the last recorded spike. This pre-processing step only had a significant impact on afferent 1.

#### Human Data

We obtained human afferent data from Dimitriou (58). In brief, human afferent data were recorded using microneurography during a visuomotor perturbation task involving 11 participants. In this task, subjects performed center-out reaching movements while a 45% counter-clockwise rotation of the visual cursor, representing the hand position, was applied.

The data included spindle activity of Ia afferents recorded from the right wrist extensor muscles and hand kinematics (wrist flexion/extension and ulnar deviation angles). Both signals were sampled at 1kHz frequency. Fascicle length was estimated as described in the “Estimating the Muscle Fascicle Stretch from Whole Muscle Stretch” section below. The fascicle length was smoothed using a moving average window (5 ms), and the afferent firing rate was smoothed using a 1-D Gaussian filter with a standard deviation of *σ* = 5 ms. Muscle velocity and acceleration were computed from the extracted muscle length by calculating the first and second derivatives.

The dataset comprised four conditions: baseline, early exposure, late exposure, and washout. Each condition included 24 trials, each 1400 ms in length, totaling 96 trials per afferent. We split the data into training, validation, and test sets using a 70/10/20 split, ensuring that the distribution of conditions was consistent across each data set.

### Estimating the muscle fascicle stretch from the whole muscle stretch

For the three datasets with recordings of Ia afferents, we estimate the fascicle length from experimentally recorded quantities.

In the human data, we used a musculoskeletal model to obtain the normalized fascicle length. Specifically, the wrist joint angles recorded during the human behavioral experiment were used to passively move a musculoskeletal model of the human hand (80) developed in MuJoCo (81). The wrist deviation and flexion were set while all other joint angles were kept at zero.

From MuJoCo, we retrieved the musculotendon length (*L*_*MT*_) of the extensor carpi radialis (ECR) muscle, which is the sum of the tendon length (*L*_*T*_) and muscle length (*L*_*M*_), by computing the equilibrium muscle configuration at each timestep of the trajectory. The muscle length was then scaled to estimate the fascicle length

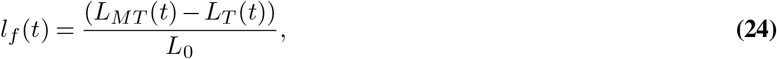

where *L*_0_ is the optimal resting muscle length. This quantity was derived using specific muscle parameters provided by MuJoCo, using the following relationship:

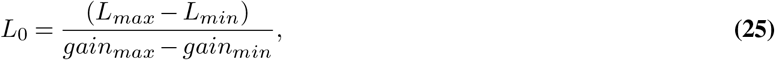

where the *gain*_*max*_ and *gain*_*min*_ are scaling factors with values of 1 and 0, respectively, and *L*_*max*_ and *L*_*min*_ are musclespecific parameters representing the maximum and minimum lengths of the muscle.

The cat and rat data had similar experimental setups (35, 55) so we used the same process to estimate the fascicle length. In both cases, a muscle puller controlled the elongation of the whole muscle length (muscle and tendon), recorded the force developed at the tendon and the length of the applied stretch. We thus had to estimate the fascicle stretch—the relevant quantity for the muscle spindle response—from the recorded applied stretch at the tendon.

The recorded length *L*_*M*_ is in units of mm and was zeroed to a resting whole muscle length at which the tension was low. We first assumed the tendon length was constant throughout the stretch and computed the normalized fascicle length from the recorded length as follows:

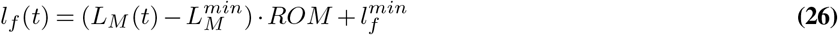

where *l*_*f*_ is the fascicle length, *L*_*MT*_ is the recorded length (in mm), 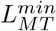 is the minimum recorded length, 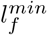 is the estimated corresponding minimum fascicle length and *ROM* (in mm^−1^) is the range of motion that we estimate as follows:

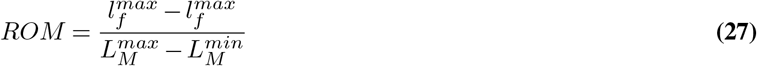

where 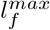 is the estimated maximal fascicle length and 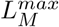 is the recorded maximal length. This process results in a fascicle length with the same profile as the recorded length quantity but with a range between 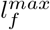 and 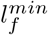. For the cat data we set 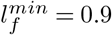 and 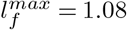. For the cat data we set 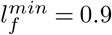 and 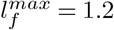.

Importantly, given that we train our models on experimental data, and that the tension equation does not depend on the fascicle length, only the fascicle velocity, a small constant offset in the fascicle length should not affect the validity of our results.

## Acknowledgments

AM is grateful to George Karniadakis for discussing PINN. We thank Mackenzie Mathis, as well as Merkourios Simos and other members of the Mathis Group for helpful feedback.

## Author contributions

A.P.R. and A.M. conceived SPIN with feedback from M.D. A.P.R. wrote the code, analyzed the data and made all figures. M.D. contributed data; M.D. and A.M.V. contributed to pre-processing of human data. A.P.R. and A.M. wrote the manuscript with input from all authors. A.M. supervised the project and acquired funding.

